# Artificial selection methods from evolutionary computing show promise for directed evolution of microbes

**DOI:** 10.1101/2022.04.01.486727

**Authors:** Alexander Lalejini, Emily Dolson, Anya E. Vostinar, Luis Zaman

**Author notes:** **Corresponding:** Alexander Lalejini.

## Abstract

Directed microbial evolution harnesses evolutionary processes in the laboratory to construct microorganisms with enhanced or novel functional traits. Attempting to direct evolutionary processes for applied goals is fundamental to evolutionary computation, which harnesses the principles of Darwinian evolution as a general purpose search engine for solutions to challenging computational problems. Despite their overlapping approaches, artificial selection methods from evolutionary computing are not commonly applied to living systems in the laboratory. In this work, we ask if parent selection algorithms—procedures for choosing promising progenitors—from evolutionary computation might be useful for directing the evolution of microbial populations when selecting for multiple functional traits. To do so, we introduce an agent-based model of directed microbial evolution, which we used to evaluate how well three selection algorithms from evolutionary computing (tournament selection, lexicase selection, and non-dominated elite selection) performed relative to methods commonly used in the laboratory (elite and top-10% selection). We found that multi-objective selection techniques from evolutionary computing (lexicase and non-dominated elite) generally outperformed the commonly used directed evolution approaches when selecting for multiple traits of interest. Our results motivate ongoing work transferring these multi-objective selection procedures into the laboratory. Additionally, our findings suggest that more sophisticated artificial selection methods from evolutionary computation should also be evaluated for use in directed microbial evolution.

## 1 Introduction

Directed evolution harnesses laboratory artificial selection to generate biomolecules or organisms with desirable functional traits (Arnold, 1998; Sánchez et al., 2021). The scale and specificity of artificial selection has been revolutionized by a deeper understanding of evolutionary and molecular biology in combination with technological innovations in sequencing, data processing, laboratory techniques, and culturing devices. These advances have cultivated growing interest in directing the evolution of whole microbial communities with functions that can be harnessed in medical, biotech, and agricultural domains (Sánchez et al., 2021).

Of course, attempting to direct evolutionary processes for applied goals has not been limited to biological systems. Evolutionary computing harnesses the principles of Darwinian evolution as a general-purpose search engine to find solutions to challenging computational and engineering problems (Fogel, 2000). As in evolutionary computing, directed evolution in the laboratory begins with a library—or population—of variants (*e.g*., communities, genomes, or molecules). Variants are scored based on a phenotypic trait (or set of traits) of interest, and the variants with the “best” traits are chosen to produce the next generation. Such approaches to picking progenitors are known as elitist selection algorithms in evolutionary computing (Bäck et al., 1997). Evolutionary computing research has shown that these elitist approaches to artificial selection can be sub-optimal in complex search spaces. On their own, elitist selection schemes fail to maintain diversity, which can lead to premature convergence (Hernandez, Lalejini, & Ofria, 2021; Lehman & Stanley, 2011a), and they lack mechanisms to balance multiple objectives. Artificial selection routines (*i.e*., parent selection algorithms or selection schemes) are intensely studied in evolutionary computing, and many *in silico* selection techniques have been developed that improve the quality and diversity of evolved solutions (*e.g*., Goings et al., 2012; Goldberg, Richardson, et al., 1987; Hornby, 2006; Lehman & Stanley, 2011b; Mouret & Clune, 2015; Spector, 2012).

Given their success, we expect that artificial selection methods developed for evolutionary computing will improve the efficacy of directed microbial evolution in the laboratory, especially when simultaneously selecting for more than one trait (a common goal in evolutionary computation). However, directed microbial evolution differs from evolutionary computing in ways that may inhibit our ability to predict which techniques are most appropriate to apply in the laboratory. For example, candidate solutions in evolutionary computing are evaluated individually, resulting in high-resolution genotypic and phenotypic information that can be used for selecting parents, which are then copied, recombined, and mutated to produce offspring. In directed microbial evolution, individual-level evaluation is often intractable at the scale required for directed evolution; as such, evaluation often occurs at the population-level, and the highest performing populations are partitioned (instead of copied) to create “offspring” populations. Moreover, when traits of interest do not benefit individuals’ reproductive success, population-level artificial selection may work against individual-level selection, which increases the difficulty of steering evolution.

Here, we ask if artificial selection techniques developed for evolutionary computing might be useful for directing the evolution of microbial populations when selecting for multiple traits of interest: both for enhancing multiple traits in a single microbial strain and for producing a set diverse strains that specialize on different traits. To do so, we developed an agent-based model of directed evolution wherein we evolve populations of self-replicating computer programs that perform computation that contributes either to the phenotype of the individual or the phenotype of the population. Using our model, we evaluated how well three selection techniques from evolutionary computing (tournament, lexicase, and non-dominated elite selection) performed in a setting that mimics directed evolution on functions measurable at the population-level. Overall, we found that multi-objective selection techniques (lexicase and non-dominated elite selection) generally outperformed the selection schemes commonly applied to directed microbial evolution (elite and top-10%). In particular, our findings suggest that lexicase selection is a good candidate technique to transfer into the laboratory, especially when aiming to evolve a diverse set of specialist microbial populations. Additionally, we found population-level artificial selection can improve directed evolution outcomes even when functional traits of interest can be tied to individual-level reproductive success.

These findings lay the foundation for strengthened communication between the evolutionary computing and directed evolution communities. The evolution of biological organisms (both natural and artificial) inspired the origination of evolutionary computation, and insights from evolutionary biology are regularly applied to evolutionary computing. As evolutionary computation has immense potential as a system for studying how to control laboratory evolution, these communities are positioned to form a virtuous cycle where insights from evolutionary computing are then applied back to directing the evolution of biological organisms. With this work, we seek to strengthen this feedback loop.

## 2 Directed evolution

Humans have harnessed evolution for millennia, applying artificial selection (knowingly and unknowingly) to domesticate a variety of animals, plants, and microorganisms (Cobb et al., 2013; Driscoll et al., 2009; Hill & Caballero, 1992; Libkind et al., 2011). More recently, a deeper understanding of evolution, genetics, and molecular biology in combination with technological advances have extended the use of artificial selection beyond domestication and conventional selective breeding. For example, artificial selection has been applied to biomolecules (Beaudry & Joyce, 1992; Chen & Arnold, 1993; Esvelt et al., 2011), genetic circuits (Yokobayashi et al., 2002), microoganisms (Ratcliff et al., 2012), viruses (Burrowes et al., 2019; Maheshri et al., 2006), and whole microbial communities (Goodnight, 1990; Sánchez et al., 2021; Swenson et al., 2000). In this work, we focus on directed microbial evolution.

One approach to artificial selection is to configure organisms’ environment such that desirable traits are linked to growth or survival (referred to as “selection-based methods” (Wang et al., 2021)). In some sense, these selection-based methods passively harness artificial selection, as individuals with novel or enhanced functions of interest will tend to outcompete other conspecifics without requiring intervention beyond initial environmental manipulations. In combination with continuous culture devices, this approach to directing evolution can be used to achieve high throughput microbial directed evolution, “automatically” evaluating many variants without manual analysis (DeBenedictis et al., 2021; Toprak et al., 2012; Wang et al., 2021). For example, to study mechanisms of antibiotic resistance, researchers have employed morbidostats that continuously monitor the growth of evolving microbial populations and dynamically adjust antibiotic concentrations to maintain constant selection on further resistance (Toprak et al., 2012). However, linking desirable traits to organism survival can be challenging, requiring substantial knowledge about the organisms and the functions of interest.

Similar to conventional evolutionary algorithms, “screening-based methods” of directed evolution assess each variant individually and choose the most promising to propagate (Wang et al., 2021). Overall, screening-based methods are more versatile than selection-based methods because traits that are desirable can be directly discerned. However, screening requires more manual intervention and thus limits throughput. In addition to their generality, screening-based methods also allow practitioners to more easily balance the relative importance of multiple objectives, such as yield, seed size, drought tolerance, *et cetera* in plant breeding (Bruce et al., 2019; Cooper et al., 2014).

In this work, we investigate screening-based methods of directed microbial evolution, as many insights and techniques from evolutionary computation are directly applicable. When directing microbial evolution, screening is applied at the population (or community) level (Sánchez et al., 2021; Xie & Shou, 2021). During each cycle of directed microbial evolution, newly founded populations grow over a maturation period in which members of each population reproduce, mutate, and evolve. Next, populations are assessed, and promising populations are chosen as “parental populations” that will be partitioned into the next generation of “offspring populations”.

Screening-based artificial selection methods are analogous to parent selection algorithms or *selection schemes* in evolutionary computing. We know from evolutionary computing research that the most effective selection scheme depends on a range of factors, including the number of objectives (*e.g*., single-versus multi-objective), the form and complexity of the search space (*e.g*., smooth versus rugged), and the practitioner’s goal (*e.g*., generating a single solution versus many different solutions). Conventionally, however, screening-based methods of directing microbial evolution choose the overall “best” performing populations to propagate (*e.g*., the single best population or the top 10% (Xie et al., 2019)).

To the best of our knowledge, the more sophisticated methods of choosing progenitors from evolutionary computing have not been applied to directed evolution of microbes. However, artificial selection techniques from evolutionary computing have been applied in a range of other biological applications. For example, multi-objective evolutionary algorithms have been applied to DNA sequence design (Chaves-González, 2015; Shin et al., 2005); however, these applications are treated as computational optimization problems. A range of selection schemes from evolutionary computing have also been proposed for both biomolecule engineering (Currin et al., 2015; Handl et al., 2007) and agricultural selective breeding (especially for scenarios where genetic data can be exploited) (Ramasubramanian & Beavis, 2021). For example, using an NK landscape model, O’Hagan et al. evaluated the potential of elite selection, tournament selection, fitness sharing, and two rule-based learning selection schemes for selective breeding applications (O’Hagan et al., 2012). Inspired by genetic algorithms, island model approaches (Tanese, 1989) have been proposed for improving plant and animal breeding programs (Ramasubramanian & Beavis, 2021; Yabe et al., 2016), and Akdemir et al. applied multi-objective selection algorithms like non-dominated selection to plant and animal breeding (Akdemir et al., 2019). In each of these applications, however, artificial selection acted as screens on individuals and not whole populations; therefore, our work focuses on screening at the population-level in order to test the applicability of evolutionary computing selection algorithms as general-purpose screening methods for directed microbial evolution.

## 3 Digital Directed Evolution

Conducting directed evolution experiments in the laboratory can be slow and labor intensive, making it difficult to evaluate and tune new approaches to artificial selection *in vitro*. We could draw directly from evolutionary computing results when transferring techniques into the laboratory, but the extent to which these results would predict the efficacy (or appropriate parameterization) of a given algorithm in a laboratory setting is unclear. To address this, we developed an agent-based model of directed evolution of microbes for evaluating which techniques from evolutionary computing might be most applicable in the laboratory.

Figure 1 overviews our model of laboratory directed microbial evolution. Our model contains a population of populations (*i.e*., a “metapopulation”). Each population comprises digital organisms (self-replicating computer programs) that compete for space in a well-mixed virtual environment. Both the digital organisms and their virtual environment are inspired by those of the Avida Digital Evolution Platform (Ofria et al., 2009), which is a well-established study system for *in silico* evolution experiments (*e.g*., A. Lalejini et al., 2021; Lenski et al., 1999; Lenski et al., 2003; Zaman et al., 2014) and is a closer analog to microbial evolution than conventional evolutionary computing systems. However, we note that our model’s implementation is fully independent of Avida, as the Avida software platform does not allow for us to model laboratory setups of directed microbial evolution (as described in the previous section).

**Figure 1:**
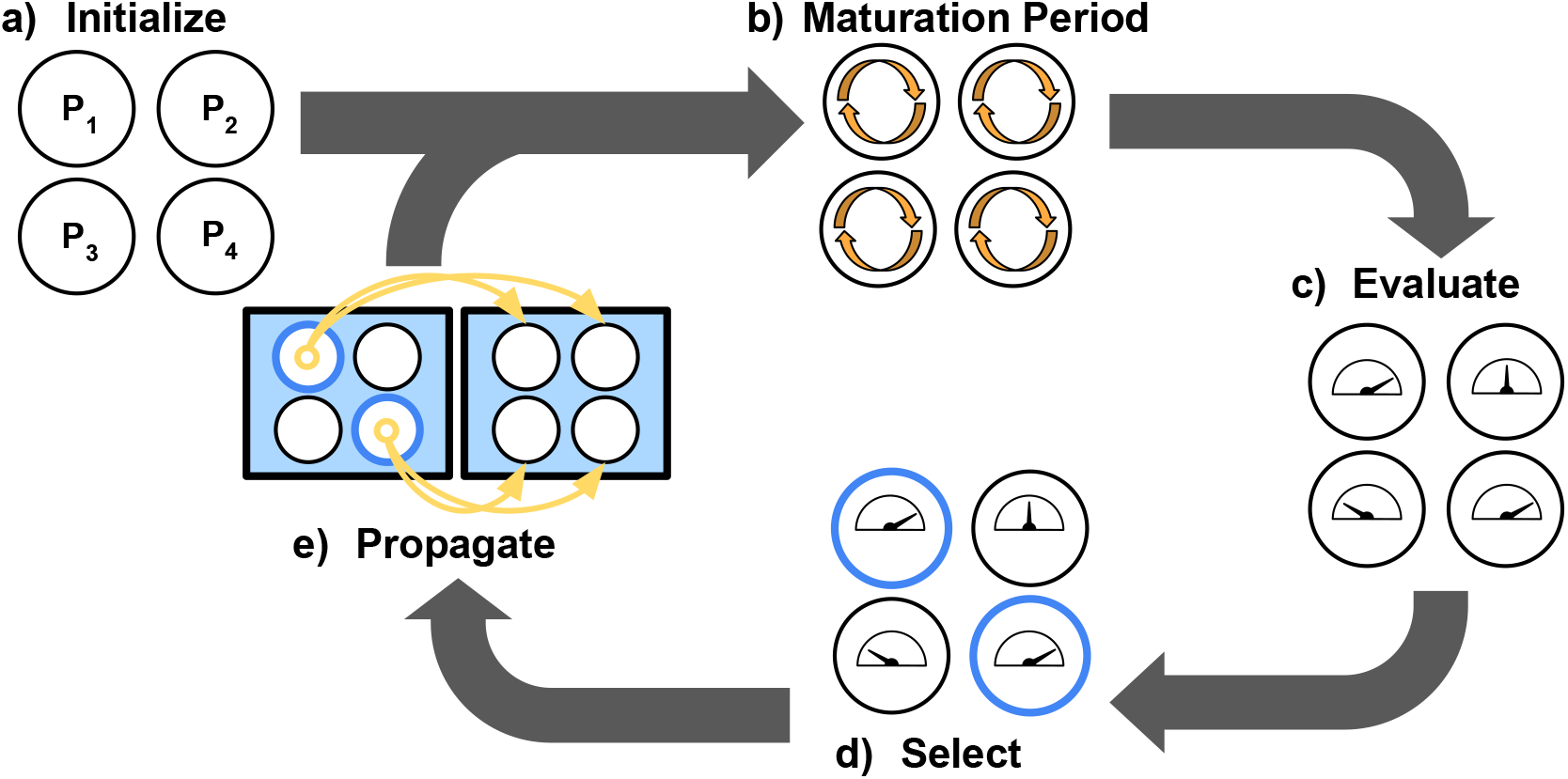
Overview of our model of directed microbial evolution. In (a), we found each of *N* populations with a single digital organism. Next (b), we give each population a maturation period during which organisms reproduce, mutate, and evolve. After maturation, (c) we evaluate each population based on population-level characteristics, and (d) we select populations (repeats allowed) to partition into N “offspring” populations (e).

In our model, we initialize each population with a digital organism capable only of self-replication (Figure 1a). After initialization, directed evolution proceeds in cycles. During a cycle, we allow each population to evolve for a “maturation period” that comprises a fixed number of time steps (Figure 1b). We then evaluate each population’s performance on a set of objectives (Figure 1c), and we select performant populations as “parental” populations (Figure 1d). To create an “offspring” population (Figure 1e), we use a random sample of digital organisms from the chosen parental population; in this work, we used 1% of the maximum population size.

### 3.1 Digital Organisms

Each digital organism is defined by a sequence of program instructions (its genome) and a set of virtual hardware components used to interpret and express those instructions. The virtual hardware and genetic representation used in this work extends that of (E. Dolson et al., 2019; Hernandez, Lalejini, & Dolson, 2021). The virtual hardware includes the following components: an instruction pointer indicating the position in the genome currently being executed, sixteen registers for performing computations, sixteen memory stacks, input and output buffers, “scopes” that facilitate modular code execution, and machinery to facilitate self-copying. For brevity, we refer readers to supplemental material for a more detailed description of these virtual hardware components (A. Lalejini et al., 2022).

Digital organisms express their genomes sequentially unless the execution of one instruction changes which instruction should be executed next (*e.g*., “if” instructions). The instruction set is Turing Complete and syntactically robust such that any ordering of instructions is valid (though not necessarily useful). The instruction set includes operators for basic math, flow control (*e.g*., conditional logic and looping), designating and triggering code modules, input, output, and self-replication. Each instruction contains three arguments, which may modify the effect of the instruction, often specifying memory locations or fixed values. We further document the instruction set in our supplemental material.

Digital organisms reproduce asexually by copying their genome instruction-by-instruction and then executing a divide instruction. However, copying is imperfect and can result in single-instruction and single-argument substitution mutations. We configured copy operations to err at an expected rate of one instruction per 100 copied and one argument per 200 copied. Genomes were fixed at a length of 100 instructions. When an organism replicates, its offspring is placed in a random position in the population, replacing any previous occupant. We limited the maximum population size to 1,000 organisms. As such, improvements to the rate of self-replication are advantageous in the competition for space within a population.

During evolution, organism replication can be improved two ways: by improving genome efficiency or by increasing the rate of genome expression (“metabolic rate”). An organism’s metabolic rate determines the speed at which it executes its genome. Digital organisms can improve their metabolic rate by performing designated functions (referred to as individual-level functions), including some Boolean logic functions and simple mathematical expressions (Table 1). Organisms can perform functions by executing the input instruction to get numeric values from the environment, performing computations on those values, and executing an output instruction with the results. When an organism produces output, we check to see if the output completes any of the designated functions (Table 1); if so, the organism’s metabolic rate is adjusted accordingly. We guarantee that the set of inputs received by an organism result in a unique output for each designated function. Organisms benefit from performing each function only once, preventing multiple rewards for repeating a single function result. In this work, we configured each function that confers an individual-level benefit to double an organism’s metabolic rate, which doubles the rate the organism can copy itself.

**Table 1:**
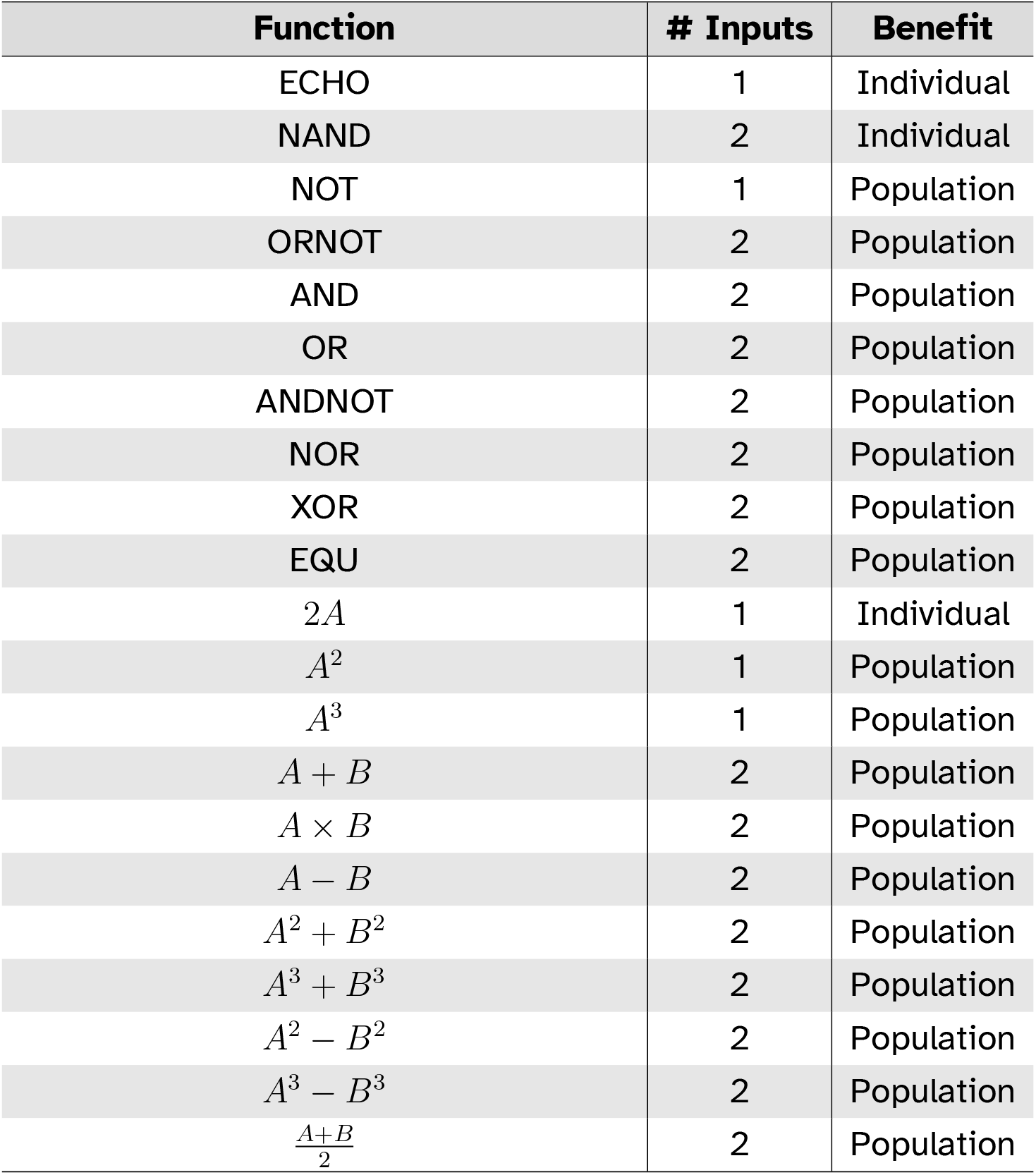
Computational functions that conferred individual-level or population-level benefits. The particular functions were chosen to be used in our model based on those used in the Avida system (Bryson et al., 2021). In all experiments, we included two versions of ECHO (each for different input values), resulting in 22 possible functions that organisms could perform.

| Function | # Inputs | Benefit |
| --- | --- | --- |
| ECHO | 1 | Individual |
| NAND | 2 | Individual |
| NOT | 1 | Population |
| ORNOT | 2 | Population |
| AND | 2 | Population |
| OR | 2 | Population |
| ANDNOT | 2 | Population |
| NOR | 2 | Population |
| XOR | 2 | Population |
| EQU | 2 | Population |
| $2A$ | 1 | Individual |
| $A^2$ | 1 | Population |
| $A^3$ | 1 | Population |
| $A + B$ | 2 | Population |
| $A \times B$ | 2 | Population |
| $A - B$ | 2 | Population |
| $A^2 + B^2$ | 2 | Population |
| $A^3 + B^3$ | 2 | Population |
| $A^2 - B^2$ | 2 | Population |
| $A^3 - B^3$ | 2 | Population |
| $\frac{A+B}{2}$ | 2 | Population |

### 3.2 Population-level Evaluation

In addition to individual-level functions, organisms can perform 18 different population-level functions (Table 1). Unless stated otherwise, performing a population-level function does not improve an organism’s metabolic rate. Instead, population-level functions are used for population-level evaluation and selection, just as we might screen for the production of different biomolecules in laboratory populations. We assigned each population a score for each population-level function based on the number of organisms that performed that function during the population’s maturation period. The use of these scores for selecting progenitors varied by selection scheme (as described in Section 4.1).

While population-level functions benefit a population’s chance to propagate, they do not benefit an individual organism’s immediate reproductive success: time spent computing population-level functions is time not spent on performing individual-level functions or self-replicating. Such conflicts between group-level and individual-level fitness are well-established in evolving systems (Simon et al., 2013; Waibel et al., 2009), and are indeed a problem recognized for screening-based methods of artificial selection that must be applied at the population-level (Brenner et al., 2008; Escalante et al., 2015).

## 4 Methods

Using our model of laboratory directed evolution, we investigated if selection schemes from evolutionary computing might be useful for directed evolution of microbes. Specifically, we compared two selection schemes used in directed evolution (elite and top-10% selection) with three other methods used in evolutionary computing (tournament, lexicase, and non-dominated elite selection). Additionally, we ran two controls that ignored population-level performance.

We conducted three independent experiments. First, we evaluated the relative performance of parent selection algorithms in a conventional evolutionary computing context, which established baseline expectations for subsequent experiments using our model of laboratory directed evolution. Next, we compared parent selection algorithms using our model of laboratory directed evolution in two contexts. In the first context, we did not link population-level functions (Table 1) to organism survival to evaluate how well each parent selection algorithm performs as a screening-based method of artificial selection. In the second context, we tested whether any of the selection schemes still improve overall directed evolution outcomes even when organism survival *is* aligned with population-level functions.

### 4.1 Selection Schemes

#### 4.1.1 Elite and top-10% selection

Elite and top-10% selection are special cases of truncation selection (Mühlenbein & Schlierkamp-Voosen, 1993) or (*μ, λ*) evolutionary strategies (Bäck et al., 1991) wherein candidates are ranked and the most performant are chosen as progenitors. We implement these selection methods as they are used in laboratory directed evolution (Xie & Shou, 2021; Xie et al., 2019). Here, both elite and top-10% selection rank populations according to their aggregate performance on all population-level functions. Elite selection chooses the single best performing population to generate the next metapopulation, and top-10% chooses the best 10% (rounded up to the nearest whole number) as parental populations.

#### 4.1.2 Tournament selection

Tournament selection is one of the most common parent selection methods in evolutionary computing. To select a parental population, *T* populations are randomly chosen from the metapopulation to form a tournament (*T* = 4 in this work). The population with the highest aggregate performance on all population-level functions wins the tournament and is chosen as a parent. We run *N* tournaments to choose the parental populations for each of N offspring populations.

#### 4.1.3 Lexicase selection

Unlike the previously described selection schemes, lexicase selection does not aggregate measures of performance across population-level functions (*i.e*., objectives) to choose parental populations. Instead, lexicase selection considers performance on each population-level function independently. For each parent selection event, all members of the metapopulation are initially considered candidates for selection. To select a parental population, the set of population-level functions are shuffled and considered in sequence. Each function (in shuffled order) is then used to sequentially filter candidates, removing all but the best candidates from further consideration. This process continues until only one candidate remains to be selected or until all functions have been considered; if more than one candidate remains, one is selected at random.

Lexicase selection was originally proposed for test-based genetic programming problems (Helmuth et al., 2015; Spector, 2012), but has since produced promising results in a variety of domains (Aenugu & Spector, 2019; La Cava et al., 2016; Metevier et al., 2019; Moore & Stanton, 2017). By randomly permuting the objectives for each parent selection, lexicase selection maintains diversity (E. L. Dolson et al., 2018; Helmuth et al., 2016), which improves search space exploration (Hernandez, Lalejini, & Ofria, 2021) and overall problem-solving success (Helmuth & Spector, 2015; Hernandez, Lalejini, & Dolson, 2021). In particular, lexicase selection focuses on maintaining specialists (Helmuth et al., 2019).

#### 4.1.4 Non-dominated elite selection

Non-dominated elite selection is a simple multi-objective selection algorithm that chooses all populations that are not Pareto dominated by another population (Zitzler, 1999). A candidate, *c_a_*, Pareto dominates another candidate, *c_b_*, if the following two conditions are met: (1) *c_a_* performs no worse than *c_b_* on all population-level functions, and (2) *c_a_* has strictly better performance than *c_b_* on at least one population-level function. After identifying all non-dominated populations, these populations are selected with replacement to found each offspring population.

Pareto domination is a fundamental component in many successful evolutionary multi-objective optimization (EMOO) algorithms (Deb et al., 2002; Fonseca & Fleming, 1995; Horn et al., 1994; Zitzler, 1999). In general, EMOO algorithms aim to produce the set of solutions with optimal trade-offs of the objective set. Most EMOO algorithms have more sophisticated routines for parent selection than non-dominated elite selection (*e.g*., use of external archives or crowding metrics). We opted to use non-dominated elite selection for its simplicity, but future work will explore more EMOO selection schemes.

#### 4.1.5 Selection controls

We used random and no selection as controls. Random selection chooses a random set of populations (with replacement) to serve as parental populations. “No selection” chooses all populations in the metapopulation as sources for founding the next generation of populations; that is, each population is chosen to produce one offspring population. Both controls apply no selection pressure for performing population-level functions.

### 4.2 Experimental design

#### 4.2.1 Establishing baseline problem-solving expectations in an evolutionary computing context

First, we evaluated the relative performance of parent selection algorithms in a conventional evolutionary computing context (linear genetic programming (Brameier & Banzhaf, 2007)), in which we evolved programs to compute the functions in Table 1. This control experiment allowed us to verify that the genetic representation used by digital organisms (Section 3.1) is sufficient for evolving each of the computational functions used in subsequent experiments. Additionally, the relative performances of each algorithm establishes an expectation for how each parent selection algorithm might perform in our model of laboratory directed evolution.

For each of the seven selection schemes described in Section 4.1, we evolved 50 replicate populations of 1,000 programs. We chose to evolve populations for 55,000 generations to approximate the number of digital organism generations that elapsed in our directed evolution experiments (based on exploratory runs). We used the same genetic representation as described in Section 3.1; however, we excluded self-replication instructions from the instruction set, as we did not require programs to copy themselves during this experiment.

Each program was evaluated independently to determine its phenotype. To evaluate a program, we executed it for 200 time steps, and we tracked its inputs and outputs to determine which of the functions in Table 1 it performed (if any). For the purpose of selection, we treated each of the 22 possible functions as a pass-fail task. Lexicase and non-dominated elite selection considered each task separately to choose parents, while elite, top-10%, and tournament selection used the number of task passes as fitness values for choosing parents. Chosen parents reproduced asexually, and we applied mutations to offspring of the same types and frequencies as in our model of laboratory directed evolution (Section 3.1).

At the end of each run, we identified the program that performed the most tasks, and we compared these values across treatments. We considered a run to be successful if it produced a program capable of performing all 22 tasks during evaluation.

#### 4.2.2 Applying parent selection algorithms in a digital directed evolution context

Next, we evaluated each selection scheme’s performance in our model of laboratory directed evolution. For each selection scheme, we ran 50 independent replicates of directed evolution for 2,000 cycles of population maturation, screening, and propagation (as shown in Figure 1). During each cycle, we gave populations a maturation period of 200 updates^1^ (approximately 25 to 35 generations). Within each replicate, the metapopulation comprised 96 populations (following the number of samples held by a standard microtiter plate used in laboratory experiments), each with a maximum carrying capacity of 1,000 digital organisms. During a population’s maturation period, we measured the number of organisms that performed each of the 18 population-level functions (Table 1) as the population’s “phenotype” for evaluation. We selected populations to propagate according to the treatment-specific selection scheme, and propagated chosen parental populations as described in Section 3.

At the end of the experiment, we analyzed the population-level functions performed by populations in each replicate’s metapopulation. First, we calculated each population’s “task profile”, which is a binary vector that describes which population-level functions are “covered” by the population (zeroes are assigned for functions that are not covered and ones for those that are covered). A function is considered covered if it is performed by at least 50 organisms (a threshold ensuring the performance was not one-off) during a given maturation period.

Next, we measured the “best population task coverage” and “metapopulation task coverage” for each replicate. Best population task coverage is measured as the number of functions covered by the population with the largest set of covered functions. Metapopulation task coverage is measured as the number of functions covered across the entire metapopulation (*i.e*., the union of unique tasks covered by each population in the metapopulation).

We also measured the phenotypic diversity within each metapopulation. Specifically, we measured the number of different task profiles present in the metapopulation (*i.e*., phenotypic richness), and we measured the “spread” of task profiles in the metapopulation. To measure a metapopulation’s task profile spread, we calculated a centroid task profile as the average of all task profiles in the metapopulation, and then we calculated the average normalized cosine distance between each population’s task profile and the centroid. A metapopulation’s task spread summarizes how different the constituent populations’ task profiles are from one another.

#### 4.2.3 Evaluating whether selection schemes improve directed evolution outcomes when population-level functions are aligned with organism survival

Selection-based methods of artificial selection tie desired traits to organism survival, eliminating the need to apply screening-based methods to populations. We tested whether the addition of population-level selection improves directed evolution outcomes even when traits of interest (population-level functions) are selected for at the individual level (*i.e*., tied to organism survival). To do so, we repeated our previously described directed evolution experiment (Section 4.2.2), except we configured all population-level functions to improve an organism’s metabolic rate in addition to the individual-level functions. As such, all population-level functions were beneficial in all treatments, including the random and no selection controls. However, only treatments with non-control selection schemes applied artificial selection at the population-level.

### 4.3 Statistical Analyses

In general, we differentiated between sample distributions using non-parametric statistical tests. For each major analysis, we first performed a Kruskal-Wallis test (Kruskal & Wallis, 1952) to determine if there were significant differences in results across treatments (significance level *α* = 0.05). If so, we applied a Wilcoxon rank-sum test (Wilcoxon, 1992) to distinguish between pairs of treatments, using a Bonferroni correction for multiple comparisons (Rice, 1989). Due to space limitations, we do not report all pairwise comparisons in our main results; however, all of our statistical results are included in our supplemental material.

### 4.4 Software and Data Availability

Our model of laboratory directed evolution is available on GitHub (see A. Lalejini et al., 2022) and is implemented using the Empirical scientific software library (Ofria et al., 2020). We conducted all statistical analyses with R version 4.04 (R Core Team, 2021), using the following R packages for data analysis and visualization: tidyverse (Wickham et al., 2019), ggplot2 (Wickham, 2016), cowplot (Wilke, 2020), viridis (Garnier, 2018), and Color Brewer (Harrower & Brewer, 2003; Neuwirth, 2014). Our source code for experiments, analyses, and visualizations is publicly available on GitHub (see A. Lalejini et al., 2022). Additionally, our experiment data are publicly archived on the Open Science Framework (see A. M. Lalejini, 2022).

## 5 Results and Discussion

### 5.1 Baseline problem-solving expectations in an evolutionary computing context

First, we established baseline performance expectations for the selection schemes in a conventional genetic programming context to validate the solvability of the individual- and population-level functions used in our digital directed evolution experiments. Two selection schemes produced successful replicates, where success is defined as evolving a program capable of performing all 22 functions: elite (1/50) and lexicase selection (47/50). No solutions evolved in any other treatment. Figure 2 depicts the number of functions performed by the best program from each replicate. All selection schemes outperformed the random and no selection controls. Differences between all pairs except random and no selection were statistically significant (Bonferroni-corrected Wilcoxon rank-sum, *p* < 0.01). Lexicase selection was the most performant followed by top-10%, elite, tournament, and non-dominated elite selection.

**Figure 2:**
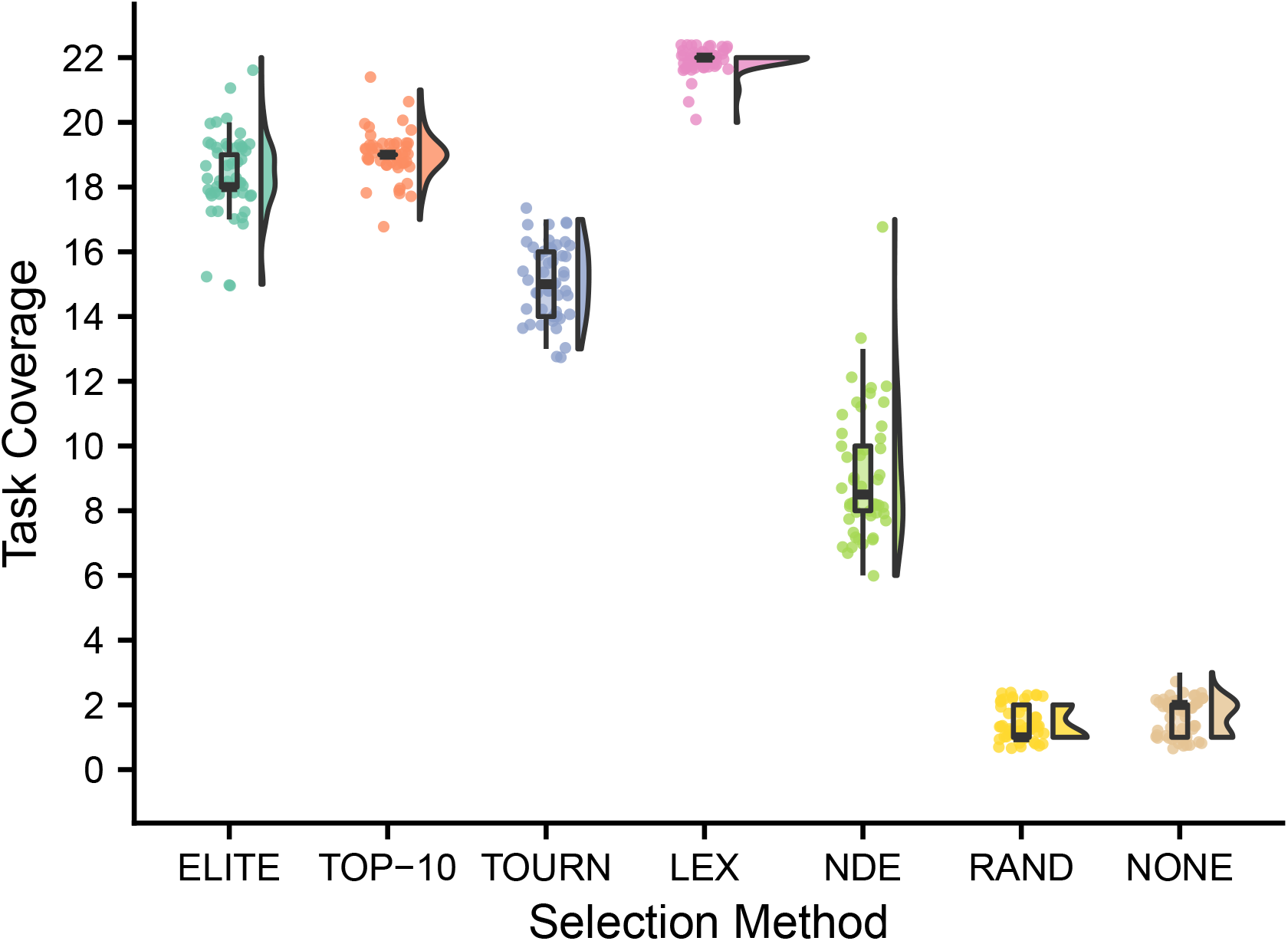
Task coverage of the best program (per replicate) evolved in an evolutionary computing context. Selection scheme abbreviations are as follows: TOURN is tournament, LEX is lexicase, NDE is non-dominated elite, RAND is random, and NONE is no selection. Differences among treatments were statistically significant (Kruskall-Wallis, *p* < 10^-4^).

These data confirm that our genetic representation allows for the evolution of each computational function used in our model of laboratory directed evolution. Moreover, these data establish some expectations for the relative performance of each selection scheme in our directed evolution experiments. Lexicase selection’s strong performance is consistent with previous work demonstrating its efficacy on program synthesis problems (Helmuth & Abdelhady, 2020; Helmuth & Spector, 2015). While initially surprised by non-dominated elite’s poor performance (relative to other non-control selection schemes), we note that selection methods based on Pareto domination are rarely applied to pass-fail test-based genetic programming problems, and perhaps the course-grained function scores (0 or 1) hindered its capacity for problem-solving success.

### 5.2 Lexicase and non-dominated elite selection show promise for directed evolution

Next, we compared selection scheme performance when modeling the directed evolution of digital organisms. Figures 3a and 3b show the best population and metapopulation task coverages, respectively. All selection schemes resulted in greater single-population task coverage than both random and no selection controls (Bonferroni-corrected Wilcoxon rank-sum test, *p* < 10^-4^). Metapopulation coverage under tournament selection was not significantly different than coverage under the no selection control, but all other selection schemes resulted in significantly better metapopulation coverage than both controls (Bonferroni-corrected Wilcoxon rank-sum, *p* < 0.03). Overall, lexicase and non-dominated elite selection scored the greatest population and metapopulation task coverage out of all selection schemes, and lexicase was the overall best selection scheme according to both metrics of performance.

**Figure 3:**
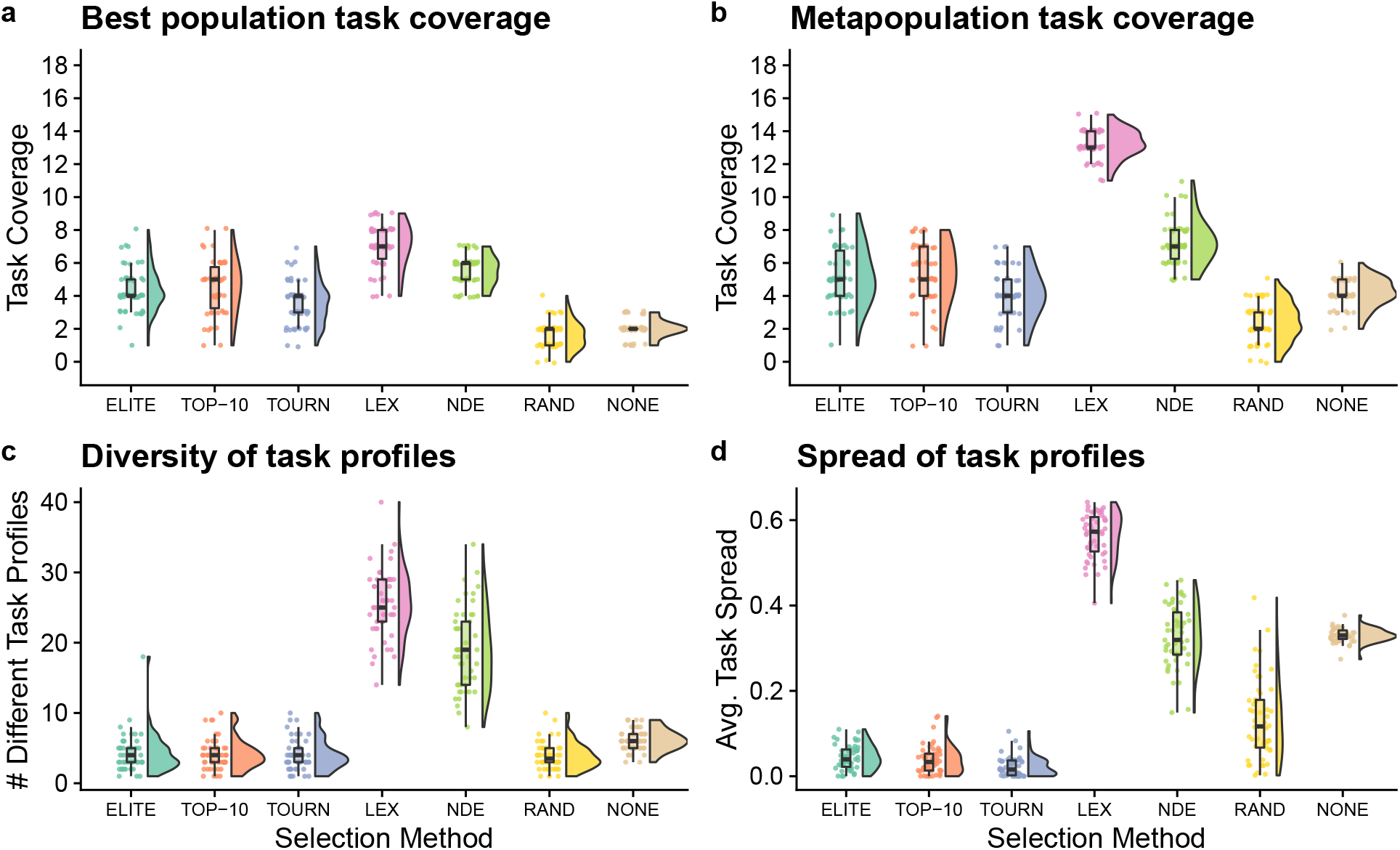
Digital directed evolution results. Differences among treatments were statistically significant for each panel (Kruskall-Wallis, *p* < 10^-4^).

While differences were significant on the best population-task coverage, they were not necessarily substantial. However, other measures had more substantial differences. Both multi-objective selection schemes—lexicase and non-dominated elite—had the greatest metapopulation task coverage (Figure 3b), and the greatest diversity of task profiles in the final metapopulations (Figure 3c; Bonferroni-corrected Wilcoxon rank-sum test, *p* < 10^-4^). Lexicase selection in particular also had the greatest task profile *spread* (Figure 3d; Bonferroni-corrected Wilcoxon rank-sum test, *p* < 10^-4^), which is consistent with previous results demonstrating that lexicase excels at maintaining diverse specialists (E. L. Dolson et al., 2018; Helmuth et al., 2016; Helmuth et al., 2019; Hernandez, Lalejini, & Ofria, 2021).

We hypothesized that lexicase and non-dominated elite selection’s mechanisms for selecting different *types* of parental populations underpinned their improved performance over elite, top-10%, and tournament selection. This, however, is confounded by each selection scheme’s varying capacity to select a greater number of different populations (regardless of differences in those selected). As such, we asked whether lexicase and non-dominated elite’s success could be explained by a capacity to select a greater number of different parental populations. Elite selection selected exactly one population per cycle, top-10% selected 10, lexicase selected an average of 12, tournament selected an average of 50, and non-dominated elite selected an average of 83 different populations. Thus, we can rule out the number of populations selected per cycle as the sole explanation for lexicase selection’s success; we argue that this, in combination with our diversity data, suggests that directed evolution practitioners should consider incorporating mechanisms for selecting phenotypically diverse parental populations into their artificial selection approaches.

These results are also informative when compared to our genetic programming control experiment (Figure 2). While results across these two contexts are not directly comparable, we argue that, taken together, our experiments suggest that steering evolution at the population-level is more challenging than steering at the individual-level. For example, across all treatments, no single population in our model of directed evolution performed all 18 population-level functions. Yet, after a similar number of organism-level generations (~55, 000), both elite and lexicase selection produced programs capable of all 22 functions in a genetic programming context; even after only 2,000 generations (the number of cycles in our directed evolution experiments), we found that conventional genetic programming produced more performant programs than those evolved under our model of laboratory directed evolution (supplemental material A. Lalejini et al., 2022). We also observed differences in the rank order of selection schemes between experiments. For example, non-dominated elite selection performed poorly in a genetic programming context relative to the other non-control selection schemes; however, non-dominated elite outperformed all selection schemes except lexicase selection in our model of laboratory directed evolution. On its own, non-dominated elite’s difference in performance is not surprising, as non-dominated elite selection is not conventionally used for evolving computer programs where evaluation criteria are evaluated on a pass-fail basis. More broadly, however, we argue that this result highlights modeling as an important intermediate step when evaluating which techniques from evolutionary computing are likely to be effective in a laboratory setting.

### 5.3 Selection schemes improve outcomes even when organism survival can be tied to population-level functions

Next, we tested whether the addition of population-level screening improves directed evolution outcomes even when population-level functions can be tied to organism survival. Overall, each non-control selection scheme resulted in better single-population task coverage than either control treatment (Figure 4a; Bonferroni-corrected Wilcoxon rank-sum test, *p* < 10^-4^). We did not find significant differences in best population coverage among elite, top-10%, tournament, and non-dominated elite selection. In contrast to our previous experiment, lexicase selection resulted in lower best population coverage than each other non-control selection scheme (Bonferroni-corrected Wilcoxon rank-sum test, *p* < 10^-4^).

**Figure 4:**
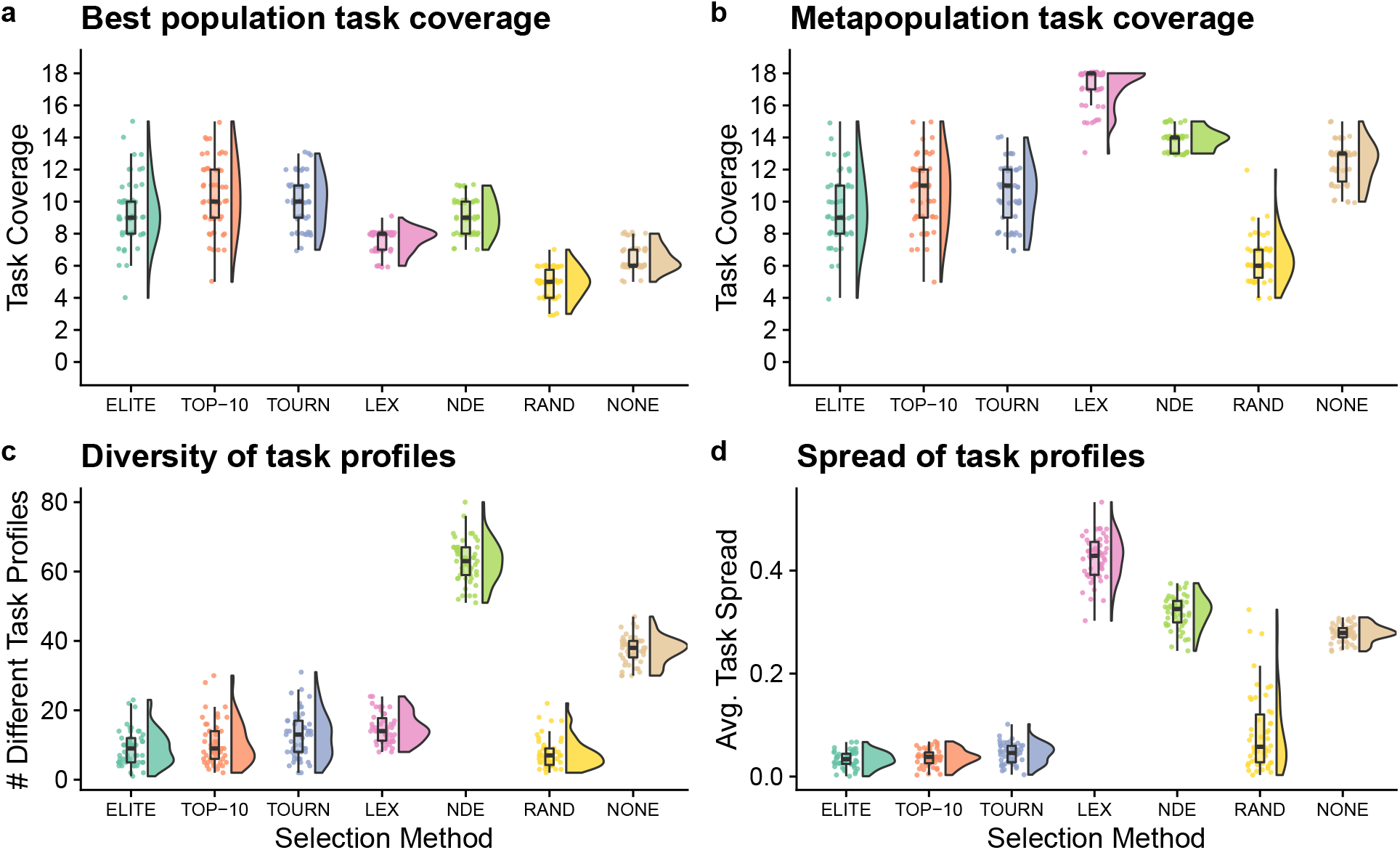
Digital directed evolution results when organism survival is tied to population-level functions. Differences among treatments were statistically significant for each panel (Kruskall-Wallis, *p* < 10^-4^).

Lexicase selection, however, outperformed all other selection schemes on metapopulation task coverage (Figure 4b; Bonferroni-corrected Wilcoxon rank-sum test, *p* < 10^-4^), producing 30 metapopulations that cover all 18 population-level functions. In general, lexicase selection produced metapopulations containing distinct specialist populations, resulting in high metapopulation task coverage while each specialist population had low task coverage on its own. Indeed, while lexicase metapopulations did not necessarily comprise many different population task profiles (Figure 4c), the task profiles were very different from one another (Figure 4d).

Of our two control selection methods, we found that performing no selection was better than random selection for both single-population and metapopulation task coverage (Bonferroni-corrected Wilcoxon rank-sum test, *p* < 10^-4^). In fact, performing no selection at all resulted in better metapopulation task coverage than elite, top-10%, and tournament selection (Bonferroni-corrected Wilcoxon rank-sum test, *p* < 10^-3^). We hypothesize that this result is because elite, top-10%, and tournament selection converge to metapopulations with homogeneous task profiles, while performing no selection at all allows populations to diverge from one another.

## 6 Conclusion

In this work, we investigated whether the selection schemes from evolutionary computing might be useful for directing the evolution of microbial populations. To do so, we introduced an agent-based model of laboratory directed evolution. Overall, our results suggest that lexicase and non-dominated elite selection are promising techniques to transfer into the laboratory when selecting for multiple traits of interest, as both of these selection schemes resulted in improved outcomes relative to conventional directed evolution selection methods. In particular, we expect lexicase selection to be especially useful for evolving a set of microbial populations, each specializing on different population-level functions. We also found that the addition of screening-based methods of artificial selection can improve directed evolution outcomes in cases where organisms’ reproductive success can be tied to traits of interest.

Our study has several important limitations that warrant future model development and experimentation. For example, we focused on modeling microbial populations that grow (and evolve) in a simple environment without complex ecological interactions. We plan to add ecological dynamics by incorporating features such as limited resources, waste by-products, symbiotic interactions, and spatial structure. These extensions will allow us to model the directed evolution of complex microbial communities (*e.g*., Sánchez et al., 2021; Xie & Shou, 2021), which is an emerging frontier in laboratory directed evolution.

In this study, we compared simple versions of each selection scheme. We plan to test more sophisticated selection schemes as we continue to transfer techniques developed for evolutionary computation into the laboratory. For example, non-dominated elite selection is one of the simplest methods that uses Pareto domination to choose parents; given its strong performance, we see more sophisticated multi-objective selection algorithms (*e.g*., NSGA-II (Deb et al., 2002)) as particularly promising for laboratory directed evolution. Lexicase selection variants are also promising for laboratory directed evolution: epsilon lexicase (La Cava et al., 2016; Spector et al., 2018) might be useful when population-level characteristics are measured as real-valued quantities, and cohort lexicase selection (Hernandez et al., 2019) could reduce the amount of screening required to select parental populations. Beyond selection schemes, we also see quality diversity algorithms (Hagg, 2021) (*e.g*., MAP-Elites (Mouret & Clune, 2015)) as promising techniques to transfer into the laboratory.

We see digital experiments like the ones reported here as a critical step for transferring techniques developed for evolutionary computing into the laboratory. Indeed, our results are currently informing the design of laboratory experiments that apply evolutionary computing techniques to the directed evolution of *E. coli*. Our model of directed microbial evolution provides a testbed for rigorously evaluating different artificial selection methods with different laboratory setups (*e.g*., metapopulation size, maturation period, *etc*.) before embarking on costly or timing consuming laboratory experiments.

## Acknowledgements

We thank the ZE^3^ Laboratory for thoughtful discussions, feedback, and support. This research was supported through computational resources provided by the University of Michigan’s Advanced Research Computing and by Michigan State University’s Institute for Cyber-Enabled Research. Additionally, this research was supported by the National Science Foundation (NSF) through support to LZ (DEB-1813069) and a sub-award to AEV (MCB-1750125).

## Supplementary information

The supplemental material for this article, including all source code, is archived (see reference) and hosted on GitHub and can be found online at https://github.com/amlalejini/directed-digital-evolution (A. Lalejini et al., 2022). The datasets generated and analyzed for this study are archived on the Open Science Framework at https://osf.io/zn63x (A. M. Lalejini, 2022).

## Footnotes

1 One update is the amount of time required for the average organism in a population to execute 30 instructions.

